# Uncovering biological patterns across studies through automated large-scale reanalyses of public transcriptomic data

**DOI:** 10.1101/2025.11.04.686647

**Authors:** Kevin Guangxing Chen, Timo Lassmann

## Abstract

Large amounts of transcriptomic data have been made available in public repositories. Systematic reanalyses of these data offer the potential to identifying conserved biological patterns or context-specific signatures. However, this is a labour intensive process requiring bioinformatic expertise and a long chain of manual decision making. Use of LLMs and agentic systems holds promise for automating these otherwise time-consuming tasks.

Here, we present UORCA (Unified -Omics Reference Corpus of Analyses), a tool to systematically identify and analyse public transcriptomic datasets. UORCA uses an LLM-assisted framework to search for datasets relevant to a research question. These datasets are analysed through a multi-agent system that performs a standardised bioinformatic analyses to identify differentially expressed genes. Results of each analysis are then displayed in an interactive visual interface.

We found that UORCA recapitulated findings reported from a manual comparison of datasets, but also found biological signatures that were not initially described. We find that UORCA generates targeted hypotheses relevant for drug design, and facilitates evaluation of experimental results where they differ from past literature. Together, these findings demonstrate how UORCA accelerates biomedical discovery by enabling scientists to extract actionable findings from diverse public datasets.

## Introduction

RNA sequencing (RNAseq)^1^ has been critical in advancing our understanding of cancer research^2^, infectious disease^3^, rare diseases^4^, and other biomedical domains. Each RNAseq dataset is generated with a specific objective in mind. However, the resulting data offer the potential to answer research questions beyond its original scope. In line with reproducibility and transparency standards, raw and processed data are routinely deposited in public repositories such as NCBI GEO, which currently encompasses over 200 000 studies^5^. Systematic reanalyses of these data therefore offer an opportunity to compare findings across biological conditions, uncovering shared and context-specific expression patterns that individual studies would not reveal.

Public data are critical across biomedical research workflows. Systematic assessments of these data, for example through literature reviews and meta-analyses, drive the generation of evidence-based hypotheses. By aggregating findings across studies, researchers can prioritise exploring questions which are supported by conserved biological signals, while also identifying unique signatures that may be indicative of novel mechanisms. Researchers will also validate new results against public data, assessing consistency with past findings. In turn, this assists in determining implications of those results, for example linking findings to biological pathways or informing therapeutic strategies.

Analysis of public RNAseq data requires extensive bioinformatic and biological domain expertise in determining suitable strategies specific to each dataset. This challenge has been recognised, with initiatives such as Galaxy^6^ and nf-core^7^ making analysis pipelines accessible and easier to use. Another consideration is the volume of public data available: databases such as ARCHS4^8^ and recount3^9^ address this by collating and harmonising public RNAseq data. These resources have been used in a diverse set of biomedical contexts, for example to identify disease-associated regulatory networks or cell subtype profiles^10–12^. These underscore the potential associated with large-scale analyses of public data. Maximising the value that can be gained from public data requires manual efforts, for example when interpreting sample metadata or identifying appropriate selections of data. As such, custom approaches are needed for each individual research question, and automation has the potential to unlock this potential broadly.

The advent of Artificial Intelligence (AI) technologies such as LLMs provides an opportunity to overcome these challenges. Specifically, the capacity of LLMs to flexibly reason over and handle variable inputs^13^ allows them to automate otherwise manual tasks. LLMs can be used to power agents: systems with the capability to call user-defined tools to achieve a set goal, such as an end-to-end RNAseq analysis. Through this, agents are able to autonomously perform analyses in the same way a human would. By tracking an agent’s tool use and the intermediate outputs, it becomes possible to reproduce and validate the agent’s analysis strategy. Agentic and other LLM-assisted approaches have been used to support transcriptomic analyses. Tools such as mergen^14^ streamline data processing steps by converting natural language queries into appropriate code. CellAgent^15^ similarly facilitates an interactive approach using a multi-agent framework. Planning, executor, and evaluation agents are used, and are each assigned a toolkit specialised to support single cell and spatial transcriptomic analysis. These approaches allow for detailed data exploration, demonstrating how LLMs and agents can enrich transcriptomic analyses. The agentic system AutoBA^16^ provides an avenue to autonomously execute a full end-to-end analysis with minimal user input, requiring only input files and the requested task. Combined, these demonstrate the capacity of using LLM-powered systems to automate tasks that traditionally required manual efforts. Agentic workflows new possibilities for systematic reanalyses of public transcriptomic data, supporting rigorous evaluation of results and increasing confidence in any reported findings.

Here, we present UORCA: Unified -Omics Reference Corpus of Analyses. UORCA empowers researchers to capitalise on the growing abundance of publicly available transcriptomic data, utilising these to generate targeted findings. UORCA provides a framework allowing automated identification and analysis of datasets, allowing researchers to systematically explore a chosen biomedical topic. We demonstrate UORCA’s capabilities and applications in streamlining comparative analyses, informing hypothesis generation, and evaluating experimental results, each drawing on the breadth of data available in public repositories. Through this, UORCA will streamline traditionally time-consuming tasks, ultimately accelerating the biomedical research workflow.

## Methods

### Overview of UORCA

The UORCA package is comprised of three distinct components: dataset identification, data analysis, and data exploration. In brief, the user inputs a research query, and an LLM-assisted workflow is used to identify relevant datasets. These datasets are then analysed via a multi-agent workflow. The researcher can then interactively explore and interrogate findings through a local web app. This overview of UORCA is shown in Figure 1A.

**Figure 1:**
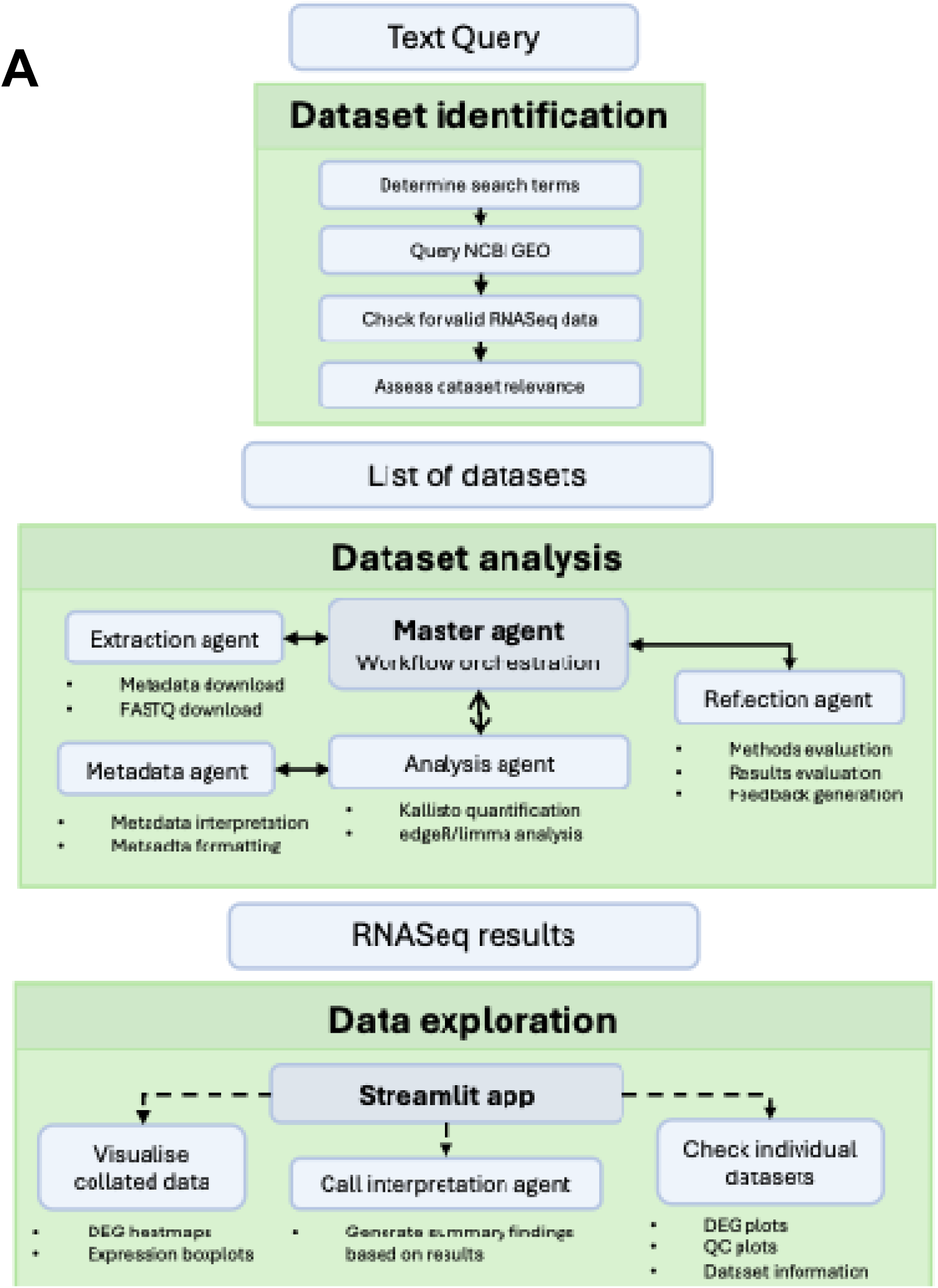
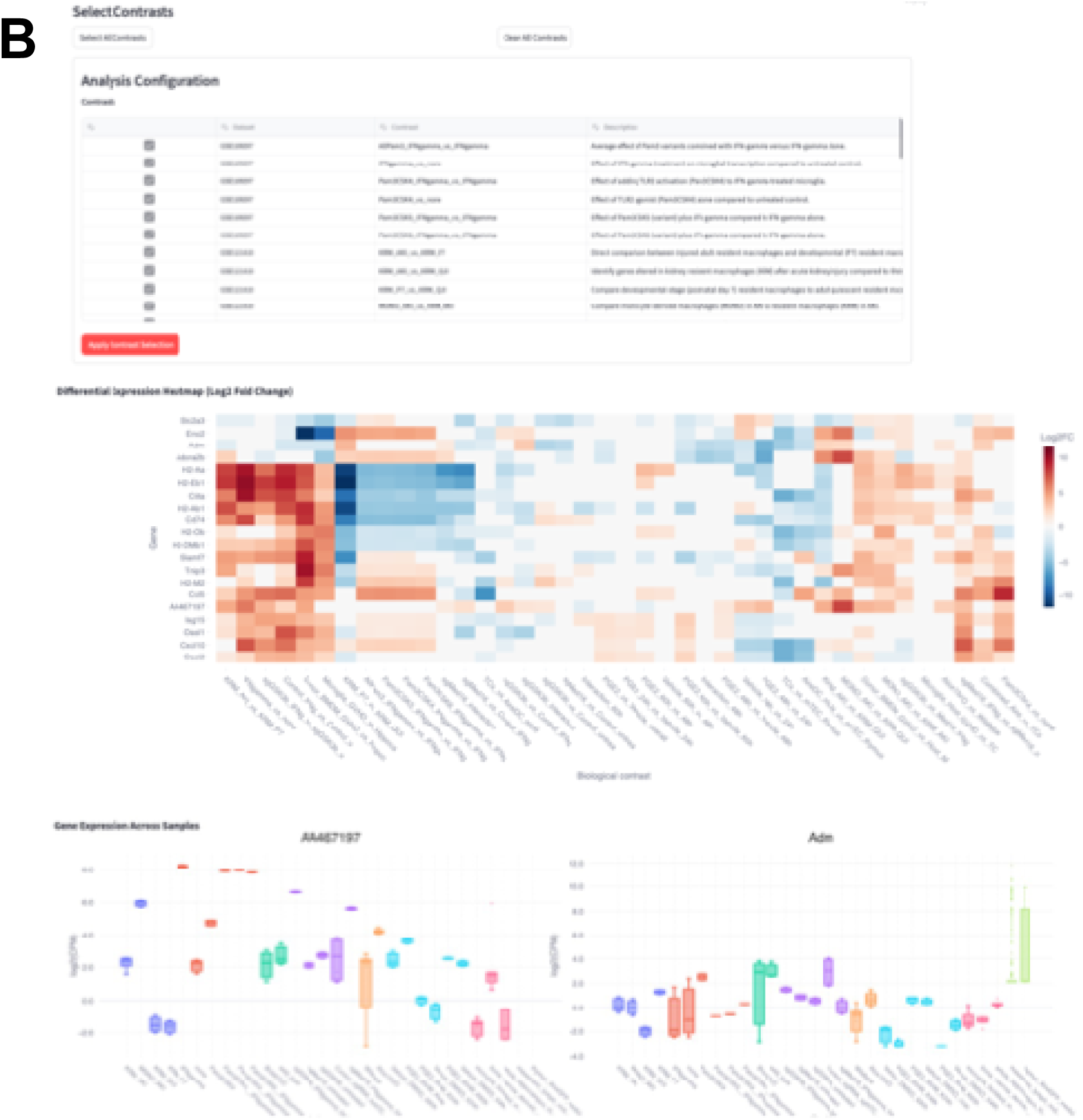
Overview of UORCA. (A) Diagram summarising the UORCA workflow. Green boxes represents the three major steps in UORCA, with the light shaded boxes showing the intermediate outputs. (B) Screenshots of the UORCA Explorer application, showing some of the available functionalities. (Top) Users are able to select which datasets and comparisons they would like to focus, (middle) for example to generate an overview of differentially expressed genes, or (bottom) view specific expression patterns.

### Dataset identification

The first step in UORCA involves identification of datasets relevant to a researcher’s query. To do so, an LLM generates search terms by extracting and expanding key words from the query. The LLM prompts can be found in Supplementary File 1. These are used for an NCBI GEO search through the ESearch function, with the “Expression profiling by high throughput sequencing” filter applied to restrict results to RNAseq datasets. After retrieving datasets, ESummary and Efetch commands are executed to extract sample and dataset metadata. The metadata are then analysed to keep bulk RNAseq datasets with paired-end reads and at least four samples; in testing, we found this to comprise the majority of datasets, and designed this first version of UORCA around these datasets to improve the system’s reliability. Datasets matching this criteria are embedded using their title and summary via OpenAI’s text-embedding-3-small model, which then undergo k-means clustering via the SKLearn package, with datasets by default clustered into groups of ten. The first member of each cluster is taken to form a set of representative datasets. An LLM call assesses these datasets for relevance to the original query based on the title and summary, assigning each a score out of ten. Datasets meeting a score threshold, by default 7, are retained to be analysed. The intermediate output can be manually validated and adjusted if desired, so users are able to adjust the datasets which are analysed in the following step.

### Data analysis agents

A multi-agent system is used to orchestrate a standard RNAseq analysis of each selected dataset. A master agent is used to coordinate and direct activity of each specialised agent: a data extraction agent, data analysis agent, and evaluation agent. In tandem, these agents execute the end-to-end analysis from download of raw files to generating differential gene expression (DGE) results.

The system prompts used for each agent can be found in Supplementary File 1.

#### Data extraction agent

To prepare the analysis stages, the raw FASTQ files, sample metadata, and dataset information must be made available. To accomplish this, an agent is assigned with two tools. The first of these tools downloads sample metadata files and stores dataset information as context. Specifically, wget is invoked to download the sample metadata file from NCBI GEO. As some datasets contain multiple types of data (e.g. RNAseq and Ampseq), this information guides the agent to download only the RNAseq files. The GEOparse package is then used to extract dataset details, including title, summary, experimental design. This information is used by other agents to inform a tailored analysis strategy. The second tool downloads the raw FASTQ files, achieved through prefetch and fasterq-dump.

#### Data analysis agent

The data downloaded by the extraction agent are used by the analysis agent to perform the RNAseq analysis. This involves quantification of FASTQ files, processing of input files to ensure they conform to required formats, interpretation of metadata, and differential gene expression analysis.

Transcript quantification is achieved via a tool calling kallisto quant, with the paired end reads and the rf-standard setting: we found these to accommodate a substantial proportion of datasets, and improved the reliability of the workflow. The agent will autonomously select an index file in this tool. By default, the index files provided are human, mouse, zebrafish, monkey, and dog. Users are able make additional files available to the agent if needed to expand the species UORCA is capable of analysing. The analysis agent is also assigned a tool to invoke a metadata agent: in summary, this agent helps inform the appropriate analysis strategy based on sample metadata and the user’s input query (see below for more details). The analysis agent then uses the DEG identification tool, using the edgeR^17,18^ and limma packages to execute a DGE analysis. This first entails gene filtering and counts normalisation via filterByExpr() and calcNormFactors() respectively. Design and contrast matrices are constructed according to the metadata agent’s analysis, and differentially expressed genes for each contrast are determined through lmFit, contrasts.fit, and eBayes.

#### Metadata analysis agent

Each dataset requires customised approaches to determine which variables are meaningful for analysis. Furthermore, as the edgeR/limma analysis has certain requirements on variable naming conventions, metadata needs to be processed to ensure compatibility. These are addressed by the metadata analysis agent, which is invoked by the analysis agent prior to DEG identification.

This agent is assigned a tool to clean metadata, involving removing special characters from column names and values. The agent also has access to a tool to combine columns, which is used to merge experimental variables into a single factor. Using this processed metadata, the agent then determines which specific biological comparisons to make, specifying the formula used to represent the comparison. To assist with this, the metadata analysis agent is given the original research question given during dataset identification via the initial invocation.

#### Reflection agent and supporting mechanisms

To improve the robustness of UORCA, a reflection agent and self-correcting mechanisms within tools are implemented to assist the agentic workflow when an error occurs.

Tools are designed to return customised traceback messages if the tool was not executed to completion. For example, if the analysis agent attempts to perform the DEG quantification without the contrasts being identified, the error message “No contrasts defined. Please run process_metadata_with_agent first” will appear. These are designed to help each agent self-correct the error.

Where self-correction is unsuccessful, a reflection agent is invoked to identify the cause of the error and plan an appropriate corrective measure. The reflection agent is given information about execution logs, previous reflections, and other information from earlier attempts. This is used by the agent to generate a plan outlining the cause of error and suggested modification. The plan is incorporated in subsequent attempts via updated prompts to agents.

#### Master agent

The master agent coordinates the overall workflow by invoking the extraction, analysis, and reflection agents. The master agent also has a tool that allows it to evaluate the analysis progress. Specifically, there are a set of checkpoints defined: FASTQ file download, metadata analysis, Kallisto index selection, Kallisto quantification, preparation for RNAseq analysis, and DEG analysis. At each checkpoint, the input parameters used as well as the output are recorded. The evaluation step checks whether analysis at each stage was successful, and also checking the parameters used to assess if they were reasonable given the available information: for example, that the Kallisto index chosen was of the correct species.

### Results visualisation

To enable users to visualise the outputs and outcomes of the UORCA-driven analyses, we include a Streamlit application that will allow researchers to interactively explore the results of the analyses. The application contains functionality to visualise common differentially expressed genes across identified contrasts, assess expression patterns for user-specified genes across all experimental groups, or view outputs from individual dataset analysis.

In the application, users can optionally use an interpretation agent which uses the analysed data to provide a summary finding in response to the user’s query. An LLM call will first determine which datasets and contrasts are most relevant to the query. The prompt used to achieve this can be found in Supplementary File 1. Following this, the log fold change data derived from these contrasts are made available to the agent, with the agent using tools to explore the data. These tools include: identifying the most common DEGs across contrasts; extracting the log fold change and p value associated with specific genes; identifying genes which are significant only in one set of contrasts, but not another; generating a summary of a specific contrast, including the number and symbols of significant genes; calculating the Spearman correlation of genes based on the average expression values; and calculating the standard deviation of log fold change values of a given gene across contrasts. For the above, the agent is able to set its own thresholds for what it deems significant. The justification of each tool is included in Table S1. The agent then outputs a set of genes and contrasts it deems relevant, and produces a summary paragraph to answer the original query.

### Output evaluation

To assess the concordance of UORCA with a typical analysis, we manually selected three datasets with differential gene expression data available. We then performed a Pearson’s correlation test comparing log fold change values in the original and UORCA analyses.

We used OpenAI’s gpt-4o-mini throughout testing of UORCA, with each individual analysis performed on a CPU cluster node (Intel Xeon Gold 6248R), using 12 cores and 16GB of memory.

## Results

### UORCA reliably analyses public RNAseq datasets

To assess the overall performance of UORCA, we ran three test cases reflecting three example applications of UORCA (Table 1). The UORCA workflow and screenshots of results visualisation are shown in Figure 1.

**Table 1:**
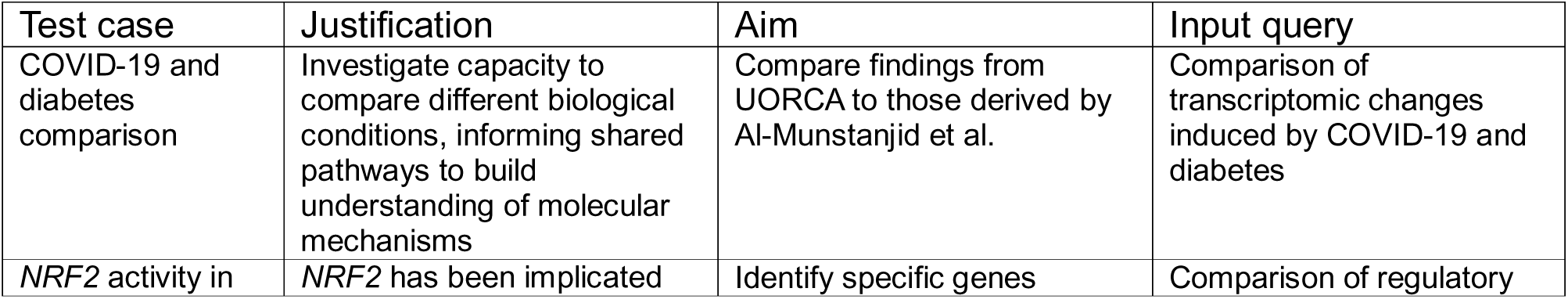
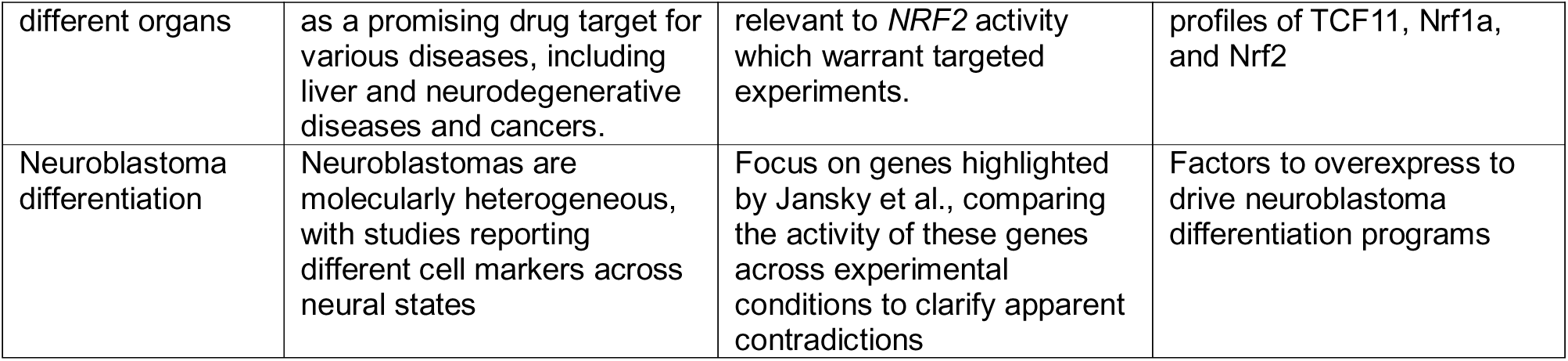
Summary of each UORCA case study.

We first aimed to evaluate UORCA’s real world performance. Across the three test cases, we found that UORCA at least 20 successful analyses in each of instance (Fig. 2), with our comparison of DGE analyses between UORCA to original findings revealing high concordance (Fig. S1). We found that the agent was able to correctly terminate analyses which could not be analysed, for example where species-appropriate files could not be found. The analysed datasets ranged from 4 -191 GB in total FASTQ file size with a median of 34 GB. Datasets took between 0.3 – 13 hours to analyse with a median of 2.2 hours (Fig. S2).

**Figure 2:**
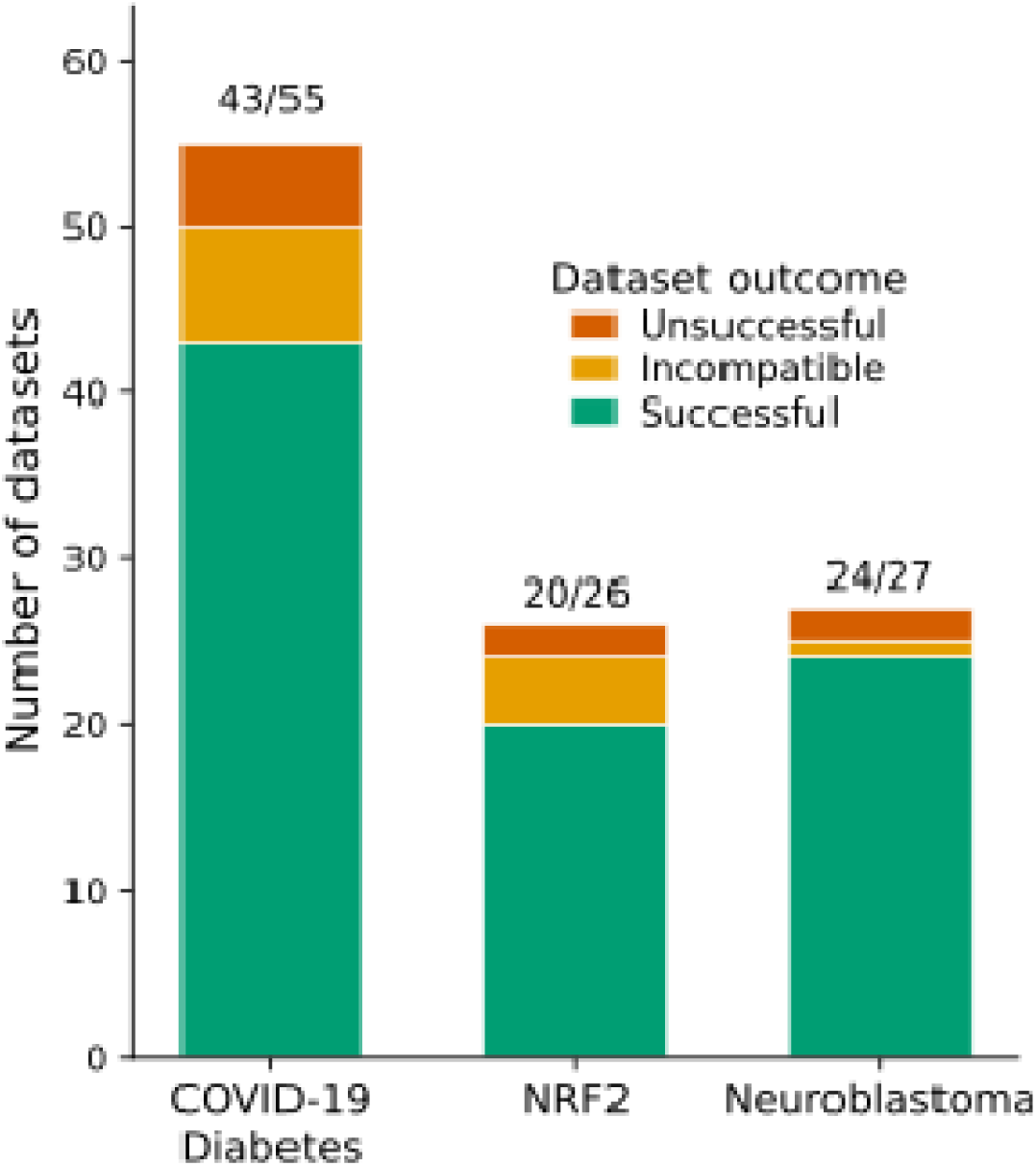
UORCA robustly analyses datasets in various biomedical contexts. Stacked barcharts showing outcomes of analyses for the three test cases, represented by each bar. The number of successful analyses out of the total analyses performed is shown on each bar.

### UORCA streamlines multi-dataset comparisons between biological contexts

We first aimed to assess the value of UORCA in analysing multiple datasets, deriving a test case from Al-Mustanjid et al.^19^ Their overall investigation examined the impact of COVID-19 on autoimmune diseases, including a comparison between a COVID-19 and Type 1 Diabetes (T1D) dataset. We emulated this with the query “Comparison of transcriptomic changes induced by COVID-19 and diabetes,” leading to automated analyses of 43 datasets. The datasets analysed by Al-Mustanjid et al. were not included in these.

The authors reported 30 shared DEGs across analysed datasets in their original study. Across datasets analysed in UORCA, three datasets (GSE201156, GSE135776, GSE160230) had at least 15 of these genes identified as differentially expressed. 29 of the 30 genes were differentially expressed in at least four datasets, including in both COVID-19 and diabetes-related datasets (Fig. 3A).

**Figure 3:**
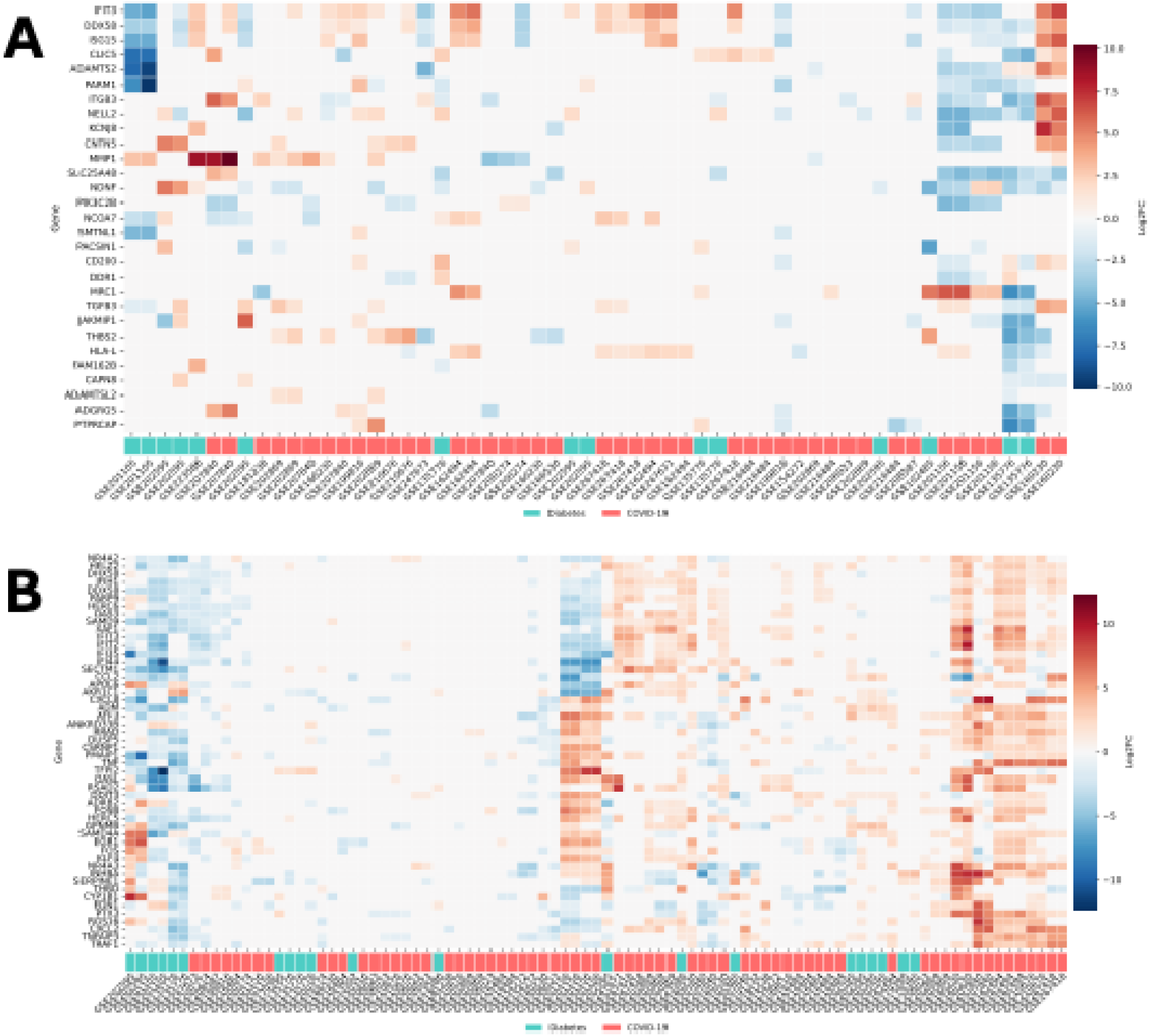
UORCA reveals shared expression patterns across diseases. Heatmap showcasing differentially expressed genes across datasets exploring diabetes and COVID-19. Heatmap colour indicates LFC as indicated by right legend. Bottom legend indicates whether the dataset was exploring COVID-19 or a diabetes. X-axis refers to the dataset that was analysed, which each column representing a specific comparison using that dataset – where a dataset is listed multiple times, this means there were multiple analyses performed using that dataset. (A) Genes highlighted by original authors, (B) top 50 most common DEGs across all analyses.

**Figure 4:**
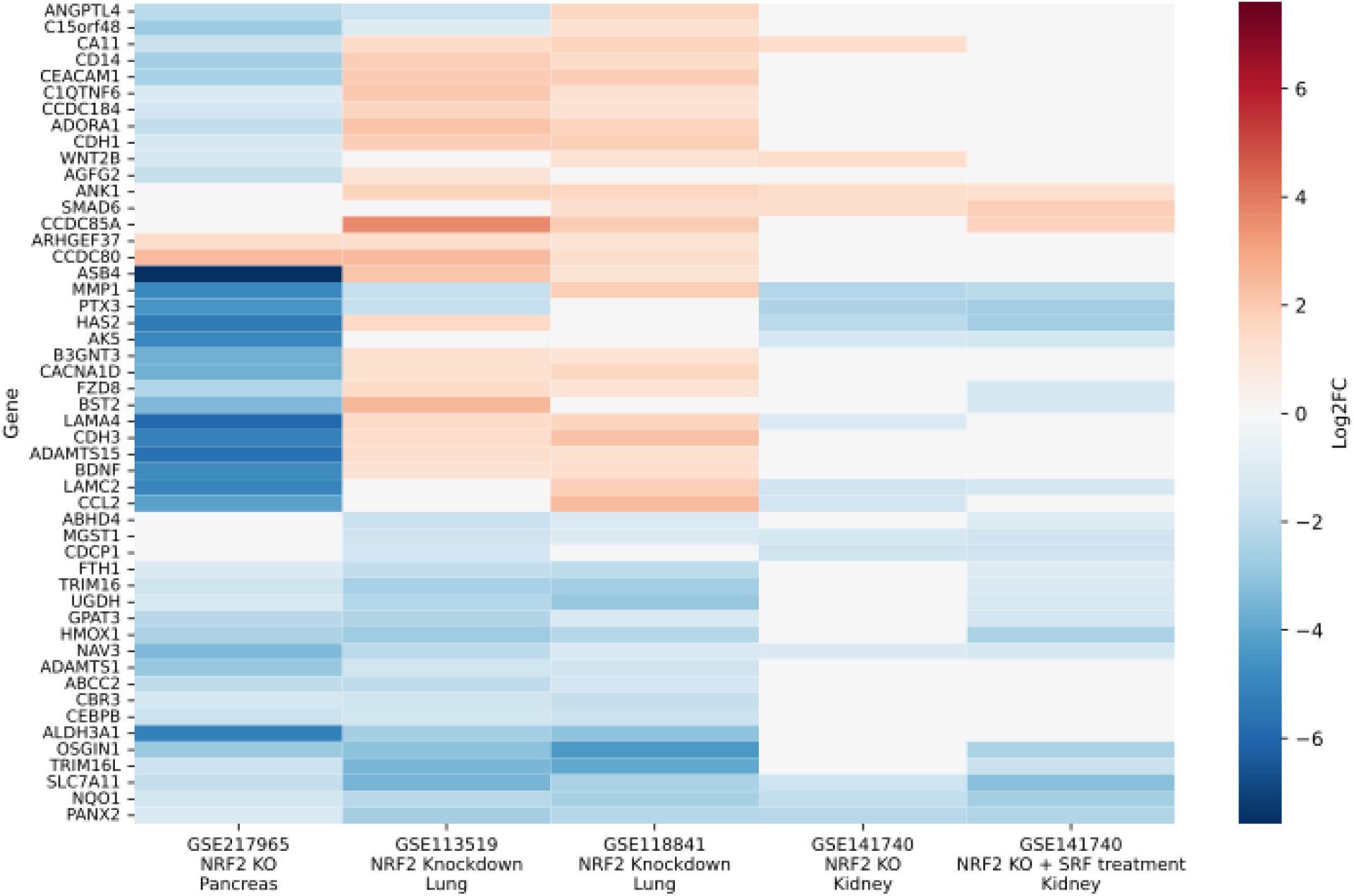
UORCA reveals tissue-specific and conserved patterns identified in datasets investigating *NRF2*. Heatmap of 50 most common DEGs across NRF2 knockout and knockdown studies. Colour indicates log fold change, with x-axis label containing basic description of each dataset, comparison, and organ.

**Figure 5:**
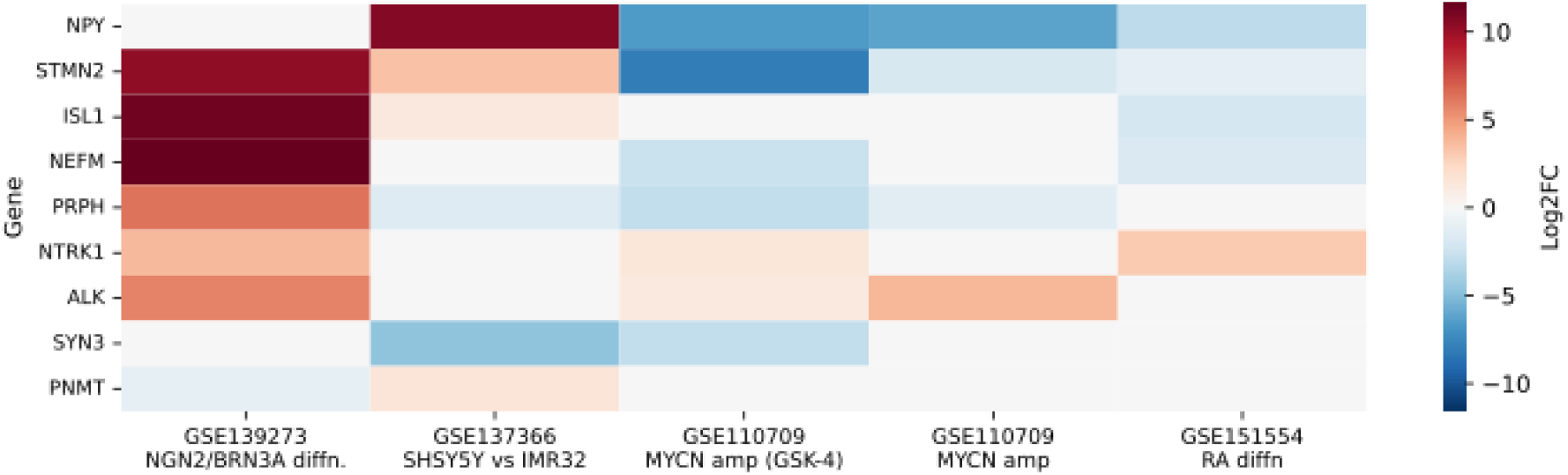
UORCA assists with explaining markers for neural cells and states. Heatmap of DEGs across neuroblastoma differentiation datasets. Colour indicates log fold change, with x-axis label containing basic description of each dataset and the specific comparison. Diffn = differentiation; amp = amplification; RA = Retinoic Acid

We next identified genes that were consistently differentially regulated across the 43 datasets (Fig. 3B). This included *IFIT3* and *DDX58*, reported by Al-Mustanjid et al. as shared DEGs between their COVID-19 and T1D dataset: these genes are involved in the interferon response, and UORCA found other interferon-induced genes as frequently differentially expressed across COVID-19 and diabetes datasets (*IFIT2, IFI35, IFI44*). Other biological themes apparent among the top 50 common DEGs included those in inflammatory responses (*CCL2*, *CXCL8*, *PTX3*) and endothelial dysfunction (*SERPINE1*, *THBD*, *EDN1*). These themes overlap with those characterised by Al-Mustanjid et al., confirming the reliability of the UORCA analyses. UORCA also revealed an apoptosis signature (*DDIT3*, *PMAIP1*, *ATF3*), which was not reported in the original findings. Apoptotic pathways have been implicated in diabetic pathologies^20,21^ and COVID-19^22,23^, demonstrating that this is a plausible biological finding. UORCA’s ability to capture this role highlights how automated analyses of many datasets enables more comprehensive results to be derived compared to what is feasible with fewer datasets.

### UORCA identifies genes with organ-specific responses to *NRF2* perturbations

*NRF2* is known to have a role in processes such as oxidative stress and inflammation, and in turn is recognised as a promising therapeutic target for neurodegenerative diseases and cancers^24–26^. As such, during an exploratory analysis we focussed on *NRF2* knockout and knockdown datasets from pancreas, lung, and kidney, identifying recurrent DEGs in human samples to highlight tissue-specific signatures. We noted some genes were consistently downregulated (*PANX2, NQO1, CEBPB*) or upregulated (*ARHGEF37, CCDC80, ANK1*) across all tissues. Other genes exhibited tissue-specific changes: *LAMC2, CDH3,* and *FZD8* were downregulated in contrasts investigating *NRF2* knockouts in kidney and pancreas cells, but upregulated in lung cancer lines. *AGFG2, WNT2B*, and *CA11* were upregulated in both kidney and lung cancer lines, but downregulated in the pancreatic cell line. Characterisation of tissue-specific differences in NRF2-regulated pathways provides insights for the design of targeted therapies. In this test case UORCA facilitated the identification of specific genes which had expression altered in response to *NRF2* perturbations. It will be of interest to study these in a focussed experiment, as findings will have implications on off-target effects if *NRF2* is to be pursued for drug development.

### UORCA clarifies cell markers across neural states

Jansky et al.^27^ performed single-cell transcriptomic analyses exploring neuroblastomas. They report *NEFM*, *STMN2*, *SYN3*, and *ALK* as neuronal markers, and PNMT as a chromaffin marker and describe contradictions pertaining to *NPY*, *PRPH*, *NTRK1*, and *ISL1* as cell markers. While Jansky et al. identified these as neuroblast-specific, these genes had been identified as chromaffin specific markers in a previous study^28^. We therefore aimed to use UORCA to clarify the nature of these markers in neuroblasts and identify biological mechanisms that may underpin results.

Exploring public datasets with UORCA and selecting these genes in the interactive application, we observed distinct expression patterns across neural cell types, developmental states, and differentiation conditions. Differentiation of patient-derived iPSCs into sensory neurons via *NGN2* and *BRN3A* overexpression^29^ led to upregulation of six neural and neuroblast markers, and downregulation of *PNMT*.

Treatment with retinoic acid (RA) or MYCN amplification all drove downregulation of *NPY* and *STMN2*. NEFM downregulation as well as upregulation of *NTRK1* were observed in RA and MYCN treatment datasets, with ALK upregulation observed in MYCN amplification analyses. These findings highlight distinct transcriptomic profiles evident at different neuroblastoma stages, with MYCN driving a proliferative undifferentiated growth, supporting the observation of the early marker *ALK*.

UORCA also identified five neural markers which were differentially expressed between the neuroblast cell lines SH-SY5Y and IMR32, despite receiving the same Nutlin treatment. This likely reflects intrinsic differences between the cell lines. The behaviour of these neural markers directly influences the conclusions which will be derived, in this case informing how to drive different neural states. As such, this highlights the need to consider how experimental results can be influenced by factors such as model or cell line choices. In sum, UORCA allows us to not just validate the findings of authors, but also contextualise these in the literature, identifying biological factors underpinning apparent contradictions.

## Discussion

Vast amounts of data have been made publicly available^30^, but tools to effectively leverage these are limited. Existing frameworks^6,15,16^ have been designed to automate narrow tasks, such as analysis of an individual study. In parallel, efforts to integrate analyses have demonstrated their value in hypothesis generation and uncovering biological relationships^31–33^. UORCA combines the capacity to automate analyses with the benefits of results integration to assist scientists with systematically exploring datasets to accelerate research.

There are concerns associated with the use of AI and LLMs in biomedical research^34–36^, namely the extent to which their output can be trusted. Agentic workflows, such as what is used in the analysis step of UORCA, perform data transformations using predefined tools. Thus, the fundamental analysis relies on well-established practices^18^, with LLMs only assisting with specific decisions. In this way, the data analyses performed by UORCA can be deterministically replicated, and the transparency offered by agentic workflows empowers researchers to manually evaluate each step of the analysis if desired.

Analysis of raw data offers opportunities not present through simple literature review. For example, the comparison made between SH-SY5Y and IMR32 cell lines was not reported by original authors^37^, highlighting how re-examining raw data can uncover findings beyond what is presented in a manuscript. UORCA autonomously processes datasets, overcoming time constraints associated with manual workflows. In doing so, UORCA facilitates more comprehensive investigations of biomedical topics compared to what is realistic with manual approaches.

Our comparison between COVID-19 and diabetes were concordant with findings reported by Al-Mustanjid et al^38^. Although there were differences in the specific genes identified as important in both COVID-19 and diabetes, this can be rationalised through dataset-specific differences; comparing alternative datasets would yield another distinct gene set^39^. More importantly, there were commonalities in the biological themes identified as important between analyses performed by UORCA and Al-Mustanjid et al, lending credence to UORCA’s value in accelerating research. In addition to the findings that were originally reported, we also identified genes that were involved in apoptosis. Comparing biological contexts between diseases helps to identify shared and context-specific pathways. In turn, this can help reveal disease mechanisms and guide drug repurposing efforts^40^.

We demonstrate the use of UORCA for hypothesis generation through an exploration of *NRF2*. *NRF2* is a well-established transcription factor, regulating a wide set of biological processes^41–43^. As a result, *NRF2* is active across many tissues. However, analyses comparing *NRF2* activity across different organs are limited^44^. As mentioned, *NRF2* has been identified as a promising drug target, and characterising off-target effects and organ-specific interactions is critical for appropriate drug development^12,45^. Even with a broad query, UORCA identified tissue-specific signatures that invite further investigation, demonstrating its utility in generating specific research questions and hypotheses.

Our evaluation of findings by Jansky et al.^27^ demonstrates how UORCA can assist researchers in contextualising results in the broader literature. Analyses performed by UORCA supported the differentiation pathway indicated by Jansky et al., and highlighted differences arising from particular growth conditions. Neuroblastomas are clinically and molecularly heterogeneous^46,47^: it is unlikely that a single dataset can appropriately capture this diversity. For diseases and conditions that display substantial diversity, it is crucial that results are comprehensively placed in context of what is known. For this purpose, UORCA not only can assist with extending experimental results to other datasets, but integrate datasets that collectively reflect disease complexity.

While UORCA is able to autonomously perform RNAseq analyses, there are still computational requirements for these analyses to be appropriately executed^48^. This is the main limiting factor determining how extensively UORCA can analyse datasets: while there is no theoretical limit on how many datasets UORCA can analyse, storage requirements and collective duration of analyses eventually become prohibitive. In the three test cases, 20, 24, and 43 datasets were explored; we found these to be sufficient, while still maintaining a reasonable time for analysis.

Overall, we demonstrate UORCA’s capacity to support researchers with automated integration of datasets. UORCA’s capabilities can be expanded by adding additional tools and agents: this will include incorporating additional repositories, data types, and analysis strategies. Moreover, the performance of UORCA will improve concordantly as LLMs become more powerful. As such, we envisage UORCA to be an evolving tool in line with growing demands of biomedical research.

### Data availability statement

The UORCA repository is available through https://github.com/Kevin-G-Chen/UORCA. The data generated for each of the three test cases can be found at https://zenodo.org/records/17402793.

## Acknowledgements

We would like to thank Timothy Chapman, Corey Ohn-Khin, and Melvin Chin for providing feedback on UORCA and this manuscript.

---

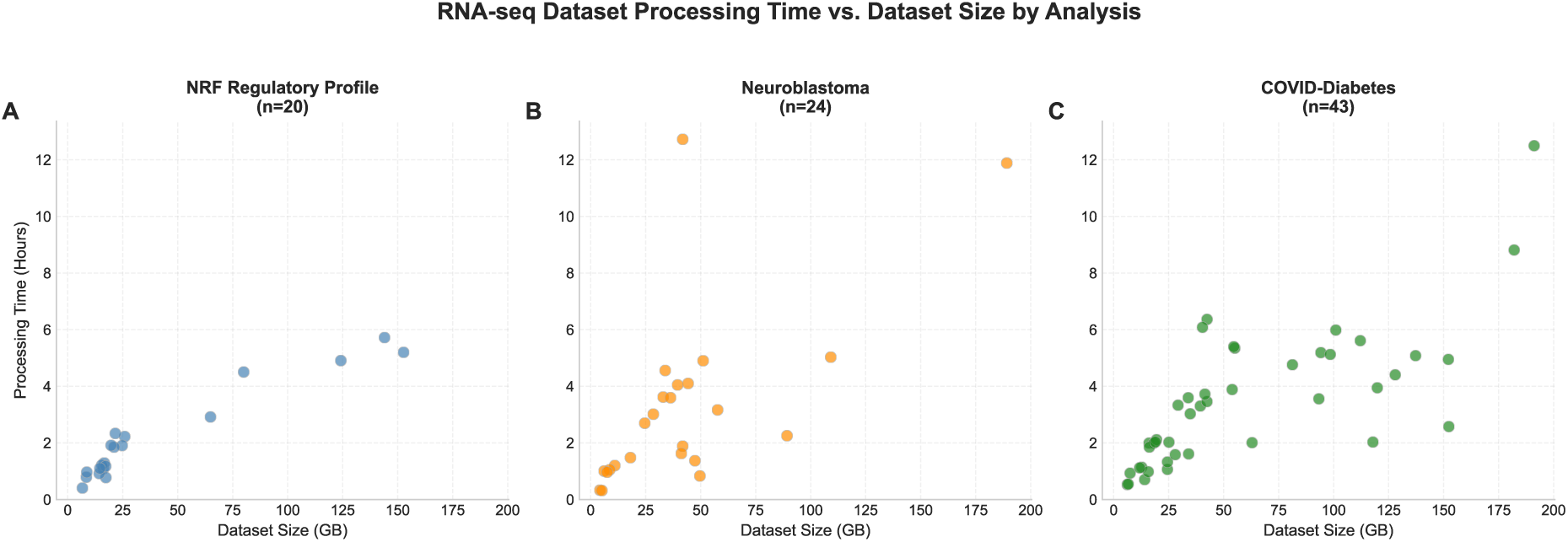

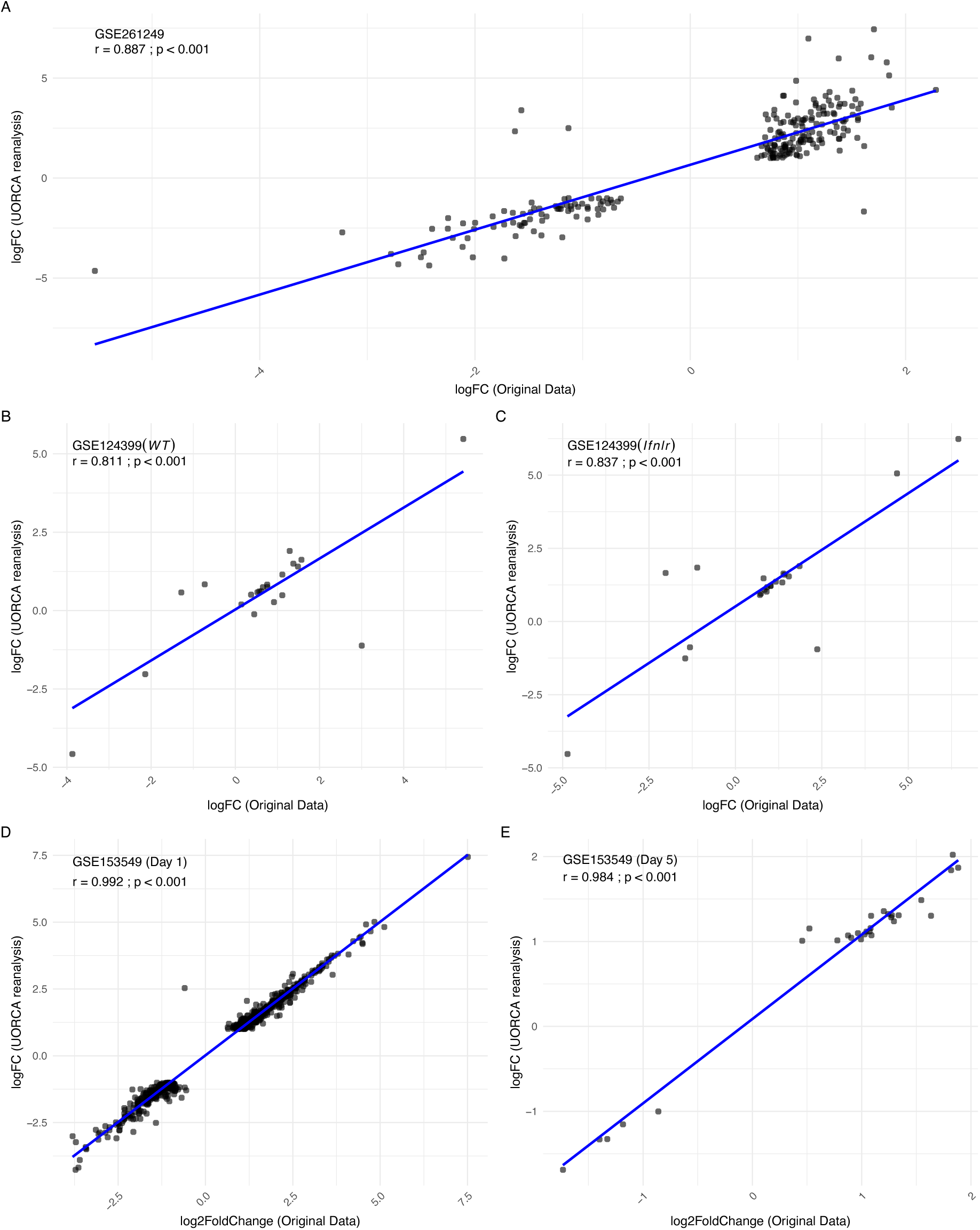

